# An Enhanced Mountain Climbing Search Algorithm to Enable Fast and Accurate Autofocusing in High Resolution Fluorescence Microscopy

**DOI:** 10.1101/2025.03.14.642633

**Authors:** Yuetong Jia, Edward N Ward, Francesca W van Tartwijk, Yutong Yuan, Yuqing Feng, Clemens F Kaminski

## Abstract

Accurate and efficient autofocusing is essential for the automation of fluorescence microscopy, but background noise and shallow depth of field at high magnifications make autofocusing particularly challenging. Here, we present a fast and accurate autofocus algorithm to address these challenges. It is highly effective for high-magnification imaging, while performing equally well for low-magnification imaging tasks. The method is based on the mountain climbing search algorithm and yields improvements on autofocusing precision of up to 200-fold over current methods, whilst offering competitive speed and greatly extended search ranges. Our approach is broadly applicable: it demonstrated good stability and reproducibility across magnifications ranging from 20X to 100X, excels in both live cell imaging and high-resolution fixed sample imaging, and it is compatible with various microscopy techniques without the need for fiducial markers or hardware modifications on existing microscopes. To maximise its accessibility, we constructed a user-friendly interface compatible with the widely used Micromanager software. It generalises well across various imaging modalities and hardware platforms, making it particularly suitable for use in high-resolution screening of candidate drugs.

## 1. Introduction

High-throughput biological screening has become increasingly important in biological and biomedical research and is used to reveal the progression of disease, targets for small molecule drugs, and mechanisms of action. [1–3] To enable higher throughput, automated and intelligent microscopy platforms have been developed to free human users from tedious and time-consuming image acquisition [4] and image processing tasks such as microscope focussing, [5] cell screening, [6] image segmentation, [7] and object classification. [8] To capture more detailed information on sample features, high-magnification objectives are needed to visualise cellular and subcellular structures. However, due to the limitation in autofocus accuracy, the automatically focused image clarity suffers as the depth of field (DOF) becomes shallower. Therefore, improvements are needed to enable more precise autofocusing during high-magnification screening.

The first automatic focusing mechanisms were based on ranging methods with sensors in 1967. [9] Since then, a variety of autofocus methods have been developed, which can be categorised into four main classes: hardware-based measurement, [10] depth information evaluation methods, [11, 12] deep learning (DL)-based autofocus, [13, 14] and iterative search approaches. [15] The hardware-based methods typically rely on reflections from an optical surface within the sample (e.g., the cover glass-sample boundary), which not only require hardware modifications like introduction of sensors, but are also difficult to implement with imaging modalities without a clear optical surface. [10] Depth information evaluation methods do not require additional hardware, improving their applicability. Instead, they evaluate the focus change across different planes to predict the defocus distance. For example, the depth from focus method [11] estimates the focus by creating a depth map with sharpness information at each point, while the depth from defocus method [12] predicts focus by calculating the degree of defocus between two images at different positions, which depends on specific lens characteristics. However, autofocus speed and stability suffer, and the methods are not easily transferable across setups due to their reliance on pre-defined models. [16] By contrast, DL approaches predict the focus directly from a single image, and thus achieve high speed. However, they require a substantial amount of data for model training and struggle to generalise models across a wide range of sample types or even sample preparation protocols. [17] Additionally, as the training data are specific to a particular optical setup, [17, 18] DL methods often fail to adapt to different experimental configurations. Because of these limitations, iterative search approaches have generally proven to be the most stable, versatile, and reliable among the four types of autofocus techniques.

Iterative search methods can be divided into three major categories: binary search, [19] rule-based search, [20] and mountain-climbing search algorithms. [15] The binary search method evaluates the image sharpness at the midpoint of the search range and then halves the search range accordingly. This iterative search continues until the sharpness value stabilises or a predefined threshold is reached. However, noise can affect the sharpness evaluation and lead to suboptimal focus. Also, this method requires the motorised stage to move back and forth to search for the optimal position, [20] with many search steps being taken, reducing the overall speed. To reduce the total iteration time, the rule-based search method has been developed, [20] which relies on pre-defined rules to adjust parameters like step size and optimise the focus. However, this requires prior knowledge of the sample and the parameters of hardware components to decide optimal rules.

The mountain climbing search method has demonstrated good performance for many automated microscopy tasks, but its accuracy and speed still pose problems for high-magnification imaging. Previous applications were shown for low magnification imaging with 10X, 20X, or 40X objectives, and several limitations were revealed. [21–23] Firstly, the search direction is typically determined by comparing the initial two or three consecutive sharpness values, but noise can lead to an incorrect search direction and autofocus failure. [22, 24] Secondly, the initial search is performed with a coarse step size to improve speed. Once a predefined sharpness threshold is reached, the step size is reduced to enhance the focus accuracy. [21, 22, 24] This threshold can vary greatly across samples, which affects the method’s general applicability. Thirdly, when imaging samples with unevenly distributed structures in three dimensions, features such as organelles might be distributed throughout the sample. As a result, each focal plane captures only a subset of these features, while structures in other planes appear blurry. This compromises the definition of “best” focus position, especially when using imaging lenses with a small DOF. Here, we address these limitations by redesigning the focus evaluation function, introducing a dynamic slope threshold, curve fitting, and an image quality evaluator, making our autofocus method broadly applicable with high accuracy.

## 2. Methods

The steps of our proposed autofocus method are shown in Fig. 1. The method uses a focus evaluation function (2.1), where the maximum defines the optimal focal plane. It consists of two distinct stages. In the first stage, the search direction is determined from a stack of five images acquired at evenly spaced planes (2.2). At the second stage, a two-step curve fitting algorithm and Naturalness Image Quality Evaluator (NIQE) are used to find the focal plane (2.3 and 2.4).

**Fig. 1.**
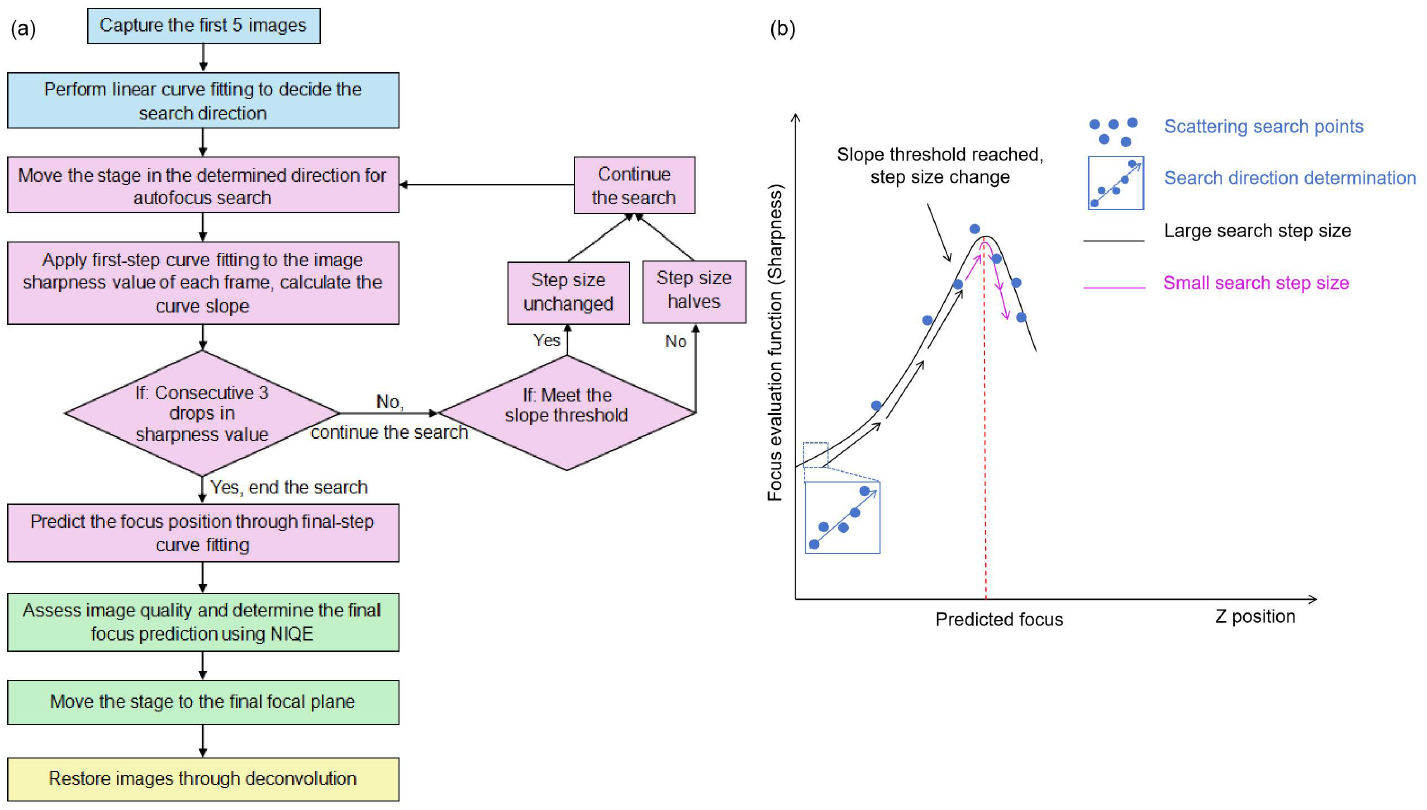
Our autofocus method follows a multi-step autofocus pipeline for high-resolution imaging. a) Workflow of the autofocus process, including four stages: initial search direction determination (blue), focus prediction using two-step curve fitting (purple), determination of the final focus position via Naturalness Image Quality Evaluator, NIQE, (green), and an optional image restoration process (yellow). b) Sketch illustrating the determination of the autofocus search direction, step adaptation, and curve fitting for focus prediction. The stage moves with the predefined search step size set by the user to capture images at various positions. The first five images (blue points) are used to determine the search direction. Then, the stage continues moving until the sharpness function’s slope meets the threshold, when the step size changes from coarse (black arrows) to fine (purple arrows). When the sharpness declines for three consecutive points, the autofocus search stops, and the optimal focus is determined from curve fitting.

### 2.1. Focus evaluation function

The focus evaluation function is the key element of the mountain climbing search algorithm, as it quantifies the image sharpness, which is in turn used to determine the search direction and define the optimal focal plane. The ideal evaluation function should exhibit a unimodal distribution within the search range and feature a clearly defined peak at the optimal focus. Additionally, the evaluation function should be applicable to different samples and be robust to noise within the image. These features are particularly important when using high-magnification imaging lenses with a shallow DOF: Here the focus evaluation function may exhibit several local maxima along the axial axis as different features within volumetric samples come in and out of focus. [25] Finally, the evaluation function needs to be computationally efficient to enable real-time evaluation. Zhang et al. [25] stated that the Laplacian gradient function is an ideal choice due to its computational efficiency, high sensitivity, and superior noise stability at high magnification (50X) compared to other evaluation functions such as Brenner and Roberts operators. However, while the Laplacian gradient shows the highest sensitivity near the optimal focus position, the sensitivity decreases at large distances from the focal plane, making it challenging to determine the focus direction if the starting position is far from optimum. An alternative focus evaluation function is the image variance operator, which shows good linearity at large distances from the focal plane, but has a lower sensitivity than the Laplacian operator closer to the in-focus plane. [26] Therefore, we combine these two operators together in our focus evaluation function.

In optical microscopy, short exposure time will limit the number of photons received by the camera, which makes background noise, including Poisson noise and Gaussian noise, more obvious. [27] Therefore, a denoising algorithm is needed to lower the influence of noise on the sharpness values.

To calculate the value of the focus evaluation function, the acquired images are first denoised through a linear background subtraction (Supplementary Notes A, Supplementary 1). The Laplacian gradient operator, ∇ ^2^ *I* (*x, y*), is then applied to the denoised images to calculate the second-order derivatives of the intensity distribution as:

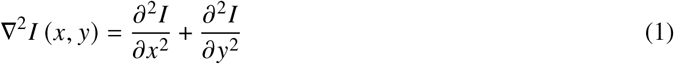

where *I x, y* is the denoised image. ∇ ^2^ *I* (*x, y*) calculates the Laplacian second-order derivative value of the input image. This detects rapid changes of intensities in the image, which indicates the presence of sharp edges.

As general intensity variations are less informative than edge variances in autofocus tasks, the edge-specific image contrast is measured by calculating the variance in the previously obtained Laplacian image using a variance operator. σ^2^ represents the variance of the Laplacian image:

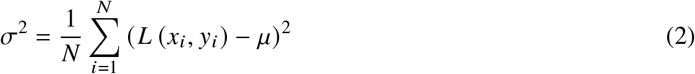

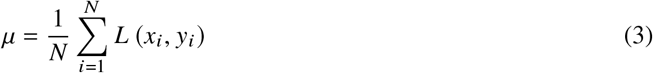

Here *N* is the total amount of pixels in an image, μ is the mean pixel value, and *L* (*x*_*i*_, *y*_*i*_) means the value of pixel (*x*_*i*_, *y*_*i*_) of the Laplacian image. The value of σ^2^ is used as the sharpness value of each image.

### 2.2. Search direction determination

The first step in the autofocus operation is to determine the search direction. In previous implementations of the mountain climbing algorithm, the search direction was determined by comparing image sharpness between two adjacent planes. [22, 24] However, since random noise increasingly affects the image sharpness at larger defocus distances, using only two adjacent frames limits the range over which this approach can function, as they may not reflect the correct trend. We therefore determine the search direction from the slope of a straight line fit through the first five search points, which greatly improves the success rate of finding the correct search direction with high NA imaging systems.

### 2.3. Two-step curve fitting with adaptive step size for focus prediction

Once the search direction has been determined, we then implement a mountain climbing algorithm to find the optimal focus. We introduce two major improvements to traditional mountain climbing methods, the first of which is the definition of search step size thresholds. Previous mountain climbing search methods required a predefined threshold value for image sharpness to guide the transition from large to small steps. [21, 22, 24] This limits general applicability: the absolute peak sharpness value varies significantly across samples and between different regions within the same sample (Fig. 2). This requires frequent adjustments of the threshold parameter, which is incompatible with unsupervised image acquisition. Instead, our algorithm autonomously determines when to switch to a smaller step size by detecting when the focus evaluation curve approaches a plateau.

**Fig. 2.**
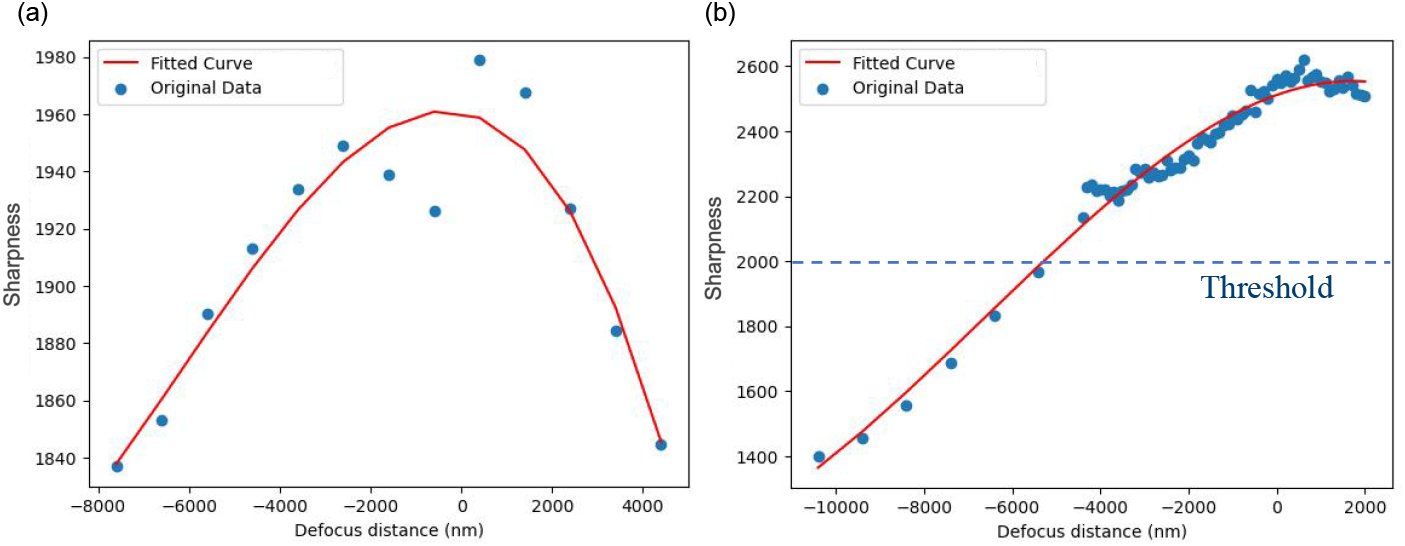
Using a predefined sharpness threshold to adjust the search step size [15] is unreliable when the field of view changes. In this example, a sharpness threshold of 2000 was set before autofocus began. a) corresponds to a field of view where the sharpness value never reaches 2000, preventing step size adaptation and leading to a poorly defined maximum. b) shows another field of view of the same sample where the threshold is reached too early before the best focus has been reached. As a result, the step size is adapted too early, prolonging the time to reach optimal focus and leading to increased photobleaching and/or phototoxicity. In both cases, the initial autofocus search parameters were: search step size of 1 μm and pre-defined sharpness value threshold of 2000. “0” in the x-axis of the plots is the optimal focus position.

To maximise the adaptability of our method, we employ both a predefined fixed gradient threshold and a dynamic gradient threshold together to detect the plateau, both of which must be met for the step size to change.

Firstly, we use a fixed slope gradient threshold, defined as:

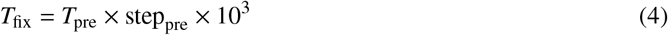

The pre-set initial search step size step_pre_ and the pre-set slope threshold *T*_pre_ are two autofocus parameters required from the user. Both the values of step_pre_ and *T*_pre_ can be determined through a single initial autofocus trial for each hardware setup. Once the optimal values are found, they remain unchanged across different regions and samples, which eliminates the need for repeated parameter adjustments between experiments. For this work, *T*_pre_ was set to 3 μ*m*^−1^ and used throughout the experiments.

In addition to these constant values, we introduced a dynamic threshold to increase the robustness of our method to noise and to account for the structural heterogeneity across different biological samples. This dynamic threshold, *T*_dyn_, is based on the average and standard deviation of the previous three slope values:

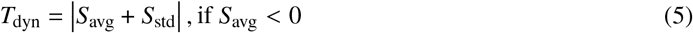

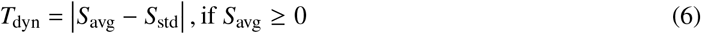

Here, *s*_avg_ represents the average value of the last three slopes, and *s*_std_ represents the standard deviation of these slope values. By evaluating the recent slope changing trend, this dynamic threshold allows for a more sensitive reflection of local changes.

In combination, the predefined threshold detects when the sharpness values start to reach a plateau, indicating a potential peak. Meanwhile, the dynamic threshold ensures that the absolute value of the slope continues to decrease rather than rise again, avoiding autofocus to stop at a local maximum. By using these two slope criteria together, we effectively avoid stationary points or saddle points, ensuring the fine search is only launched near the curve’s peak.

Our second change to traditional mountain climbing algorithms is the use of curve fitting. Traditional mountain climbing search methods require the sample stage to move back and forth in order to evaluate the focus of each frame. [22] This repeated stage movement continues until the sharpness value threshold is reached, which requires a considerable amount of time. Furthermore, the pre-defined minimum step size determines the highest precision of stage adjustment, restricting the achievable autofocus accuracy. We solve these problems through the use of a curve fitting approach, which increases the autofocus precision through mathematical model predictions and iterative optimisation. This allows the optimal focus to be determined to a higher precision than the step size and means that autofocus can be achieved in a single pass.

Here, the least square fitting method is chosen for its higher speed. In the absence of higher order aberrations (e.g. astigmatism), the image sharpness curve around the optimal focus is expected to follow a negative quadratic distribution, with a single maximum while being approximately symmetrical around the peak (See Fig. 1 (b)). This symmetric sharpness distribution is explained in Supplementary Notes B, Supplementary 1. As orthogonal polynomial functions often lead to higher stability, Legendre polynomials are used to fit the curve to the sharpness data points around the focal plane. Based on the quadratic approximation, we choose the final ten search points as the region for the curve fitting.

Our resulting autofocus procedure incorporating these changes is a two-step process that minimises stage movement and maximises accuracy. During the “first-step curve fitting”, the slope value at each position is predicted by fitting a quadratic curve with the preceding ten search points. When the slope value reaches both the fixed and the dynamic threshold for three consecutive search points, the search step size is halved to enable finer focusing. Our method allows the search step size to be halved for the second time if both criteria are met again before the search ends, allowing for more precise searches. The mountain climbing search ends when the corresponding sharpness value of the next three consecutive search points continues to decrease. After the search ends, “second-step curve fitting” launches to fit the last ten data points and calculate the fitted curve peak position as the predicted focus position.

### 2.4. NIQE focus precision enhancement

To further improve the reliability of our method, we additionally include an error-checking mechanism as the final stage of the focussing process. While the curve fitting predicts an ideal focus position, we observed that, in samples where multiple planes could be considered as in focus, the peak of the fitted curve can lie in a plane with a sub-optimal focus. To address this, the quality of the image at the predicted ideal focus (Fig. 3, point A) is compared to that of the image with the highest sharpness value that was acquired during the mountain climbing process (Fig. 3, point B). This image quality assessment is performed using a no-reference quality assessment evaluator, NIQE. [28, 29] NIQE (Fig. 4) uses a pretrained machine learning model based on human perception evaluation, and evaluates image quality based on criteria such as blur, illumination, noise, and distortion. [30] The image with the lowest NIQE score is chosen as the final output focus plane. We note that this error-checking step is generally not required for low-magnification imaging or when deconvolution (Section 2.5) is used to bring multiple adjacent planes into focus simultaneously.

**Fig. 3.**
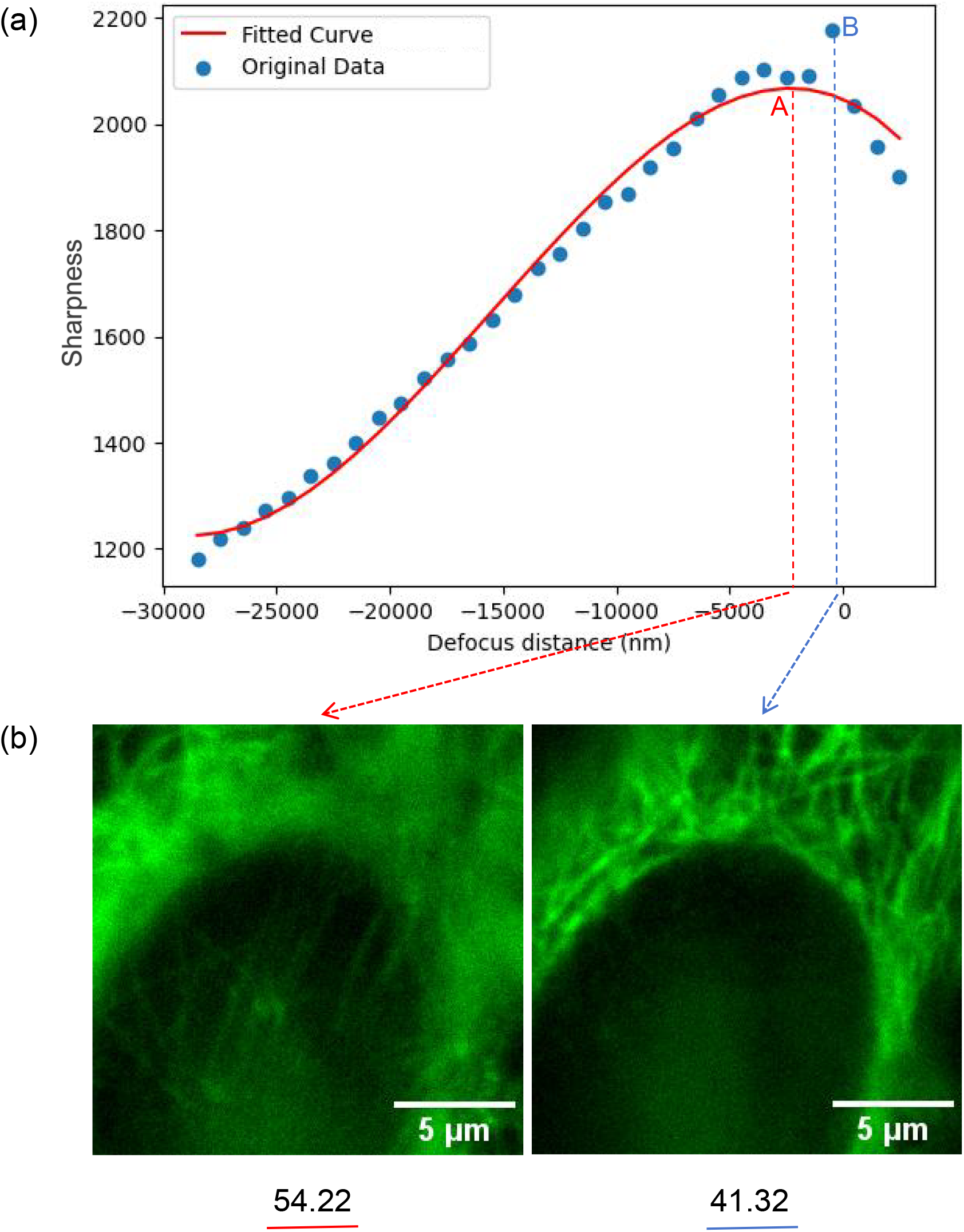
The NIQE method improves final focus position determination accuracy by assessing the image quality at two “peaks”. a) The plot shows two “peaks”: red point A, the peak position of the fitted curve, and blue point B, the scattering search data point with the highest sharpness value. The scattered blue points are the mountain climbing search data, and the red curve is the corresponding fitted curve. The peak search position (blue dashed line) is closer to the optimal focal plane (0 defocus distance) than the curve fitting predicted focus (red dashed line). b) The NIQE assessment scores for images at peak A and B show that peak B, with the better image quality (lower NIQE score), is the final and more accurate focus position. deconvolution [34, 35] to produce an output stack with improved image quality. As the brightest regions usually represent target structures and signals of interest in fluorescence imaging, a maximum intensity projection is then used to combine this stack into a single image, ensuring all the structures are in focus simultaneously in the final image.

**Fig. 4.**
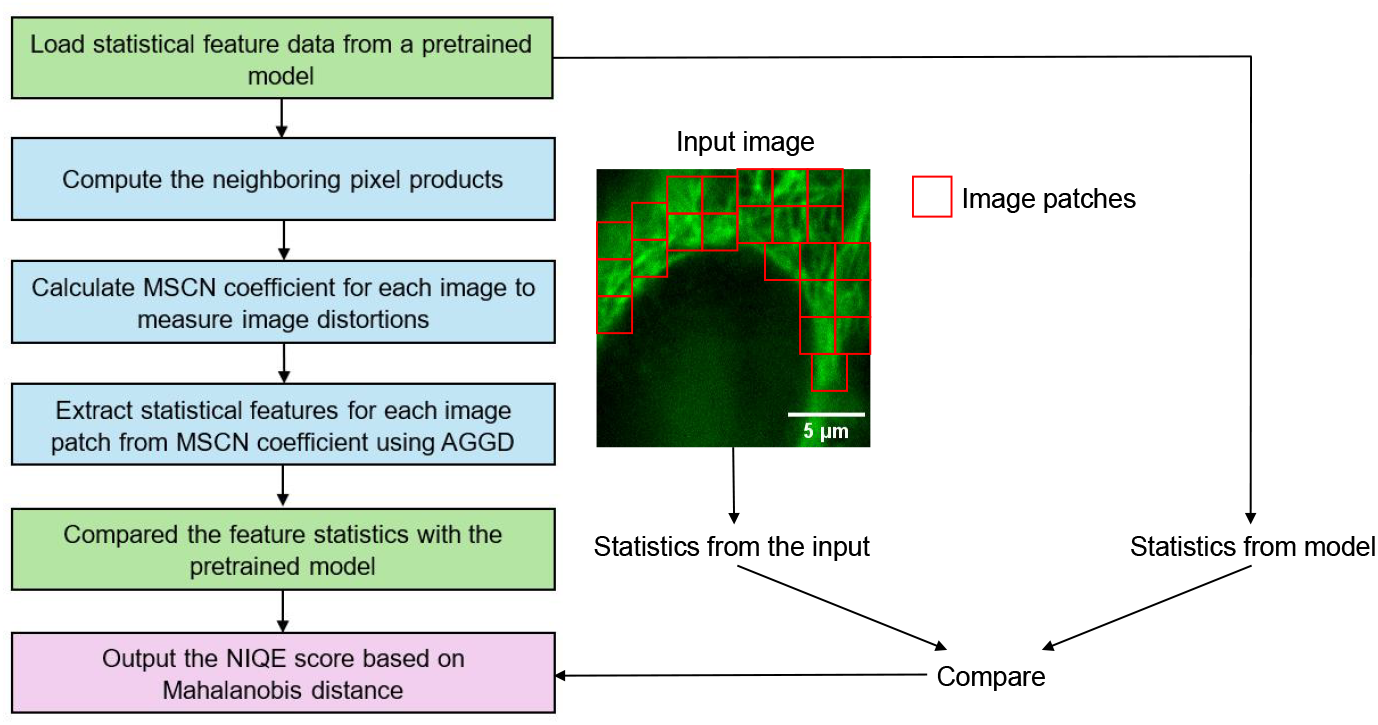
NIQE measures image quality by comparing the statistical features calculated from the input with that of a pretrained model. It can be divided into three main parts: statistical feature extraction (blue), model comparison (green), and image quality scoring (purple). Firstly, original statistical feature data is loaded from a pretrained model, which is trained using high-quality natural images. Then, as neighbouring pixels are always correlated, their products are calculated to map the spatial relationships between pixels. The statistical features Mean Subtracted Contrast Normalised (MSCN) coefficients and asymmetric generalised Gaussian distribution (AGGD) are calculated and compared with the pretrained model. The degree of deviation from the pretrained model gives an image quality evaluation score, where lower scores indicate better image quality.

### 2.5. Deconvolution using acquired z-stacks

In a final image processing step, we take advantage of the out-of-focus frames recorded during the autofocusing procedure. These data were used to deconvolve the final images, improving image quality for thicker samples with 3D distributed structures. We use the Born and Wolf theoretical point spread function (PSF) modelling method [31–33] to estimate the system PSF. The PSF model is then used to perform a 3D Richardson–Lucy (RL)

### 2.6. Graphical user interface

To make our method more accessible to non-expert users, we built a graphical user interface (GUI, Fig. S1, Supplementary 1) that is compatible with the widely used Micromanager software. [36, 37] Standard parameters like exposure time and channels are set within Micromanager, and the autofocus-specific settings such as initial search step size and the predefined slope threshold are selected through our GUI.

### 2.7. Autofocus performance quantification

Autofocus error and total focus time are used as two parameters to evaluate the performance of autofocus methods. Here, the error is quantified by calculating the deviations between the predicted focus position and the optimal focus position, which is determined manually as the position at which most of the structures within the field of view are clear. Previous studies that define optimal focus through objective measures, such as sharpness metrics and curve peak, [21, 22] are suitable for low-magnification objectives, but are unreliable when the objective magnification increases and DOF decreases: multiple focal planes within cells bring different structures into focus, meaning the highest sharpness value may not always correspond to the optimum plane for observation. Therefore, subjective optimal focal plane determination is needed for more accurate autofocus performance quantification.

The autofocus error is shown in the format of “mean ± standard error of the mean (SEM)”. The focussing time has four main components: the time it takes to determine the initial search direction, the image capture time, the time required for stage movement, and the processing time for the algorithm (including sharpness evaluation, two-step curve fitting, and NIQE comparison). Lower autofocus error and shorter focus times indicate better autofocus performance.

## 3. Results

To evaluate the precision of our autofocus method and its advantages at high magnification, we compared its performance with current state-of-the-art approaches. First, we compare performance at different defocus distances. We then demonstrate the general applicability of our method by testing it across different magnification objectives, various types of samples, and for different microscopy techniques. Descriptions of the systems we used for image acquisition and the sample preparation protocols are provided in Supplementary Notes C and Sample Preparation in Supplementary 1.

### 3.1. Our autofocus method outperforms existing methods at shallow depth of field

First, we benchmarked our method against two established autofocus methods, JAF and OughtaFocus, to demonstrate the improved performance at high magnification. Specifically, we used a 100X 1.49NA oil-immersed objective, as the high numerical aperture and magnification result in an extremely shallow DOF. Fig. 5 shows a comparison of the techniques for both autofocus accuracy and autofocus duration when imaging fixed Vero cells with immunolabelled *β*-tubulin. To show the generalisability of the method, we additionally run the experiment on a sample of live COS-7 cells with MitoTracker-stained mitochondria (Fig. S2, Supplementary 1).

**Fig. 5.**
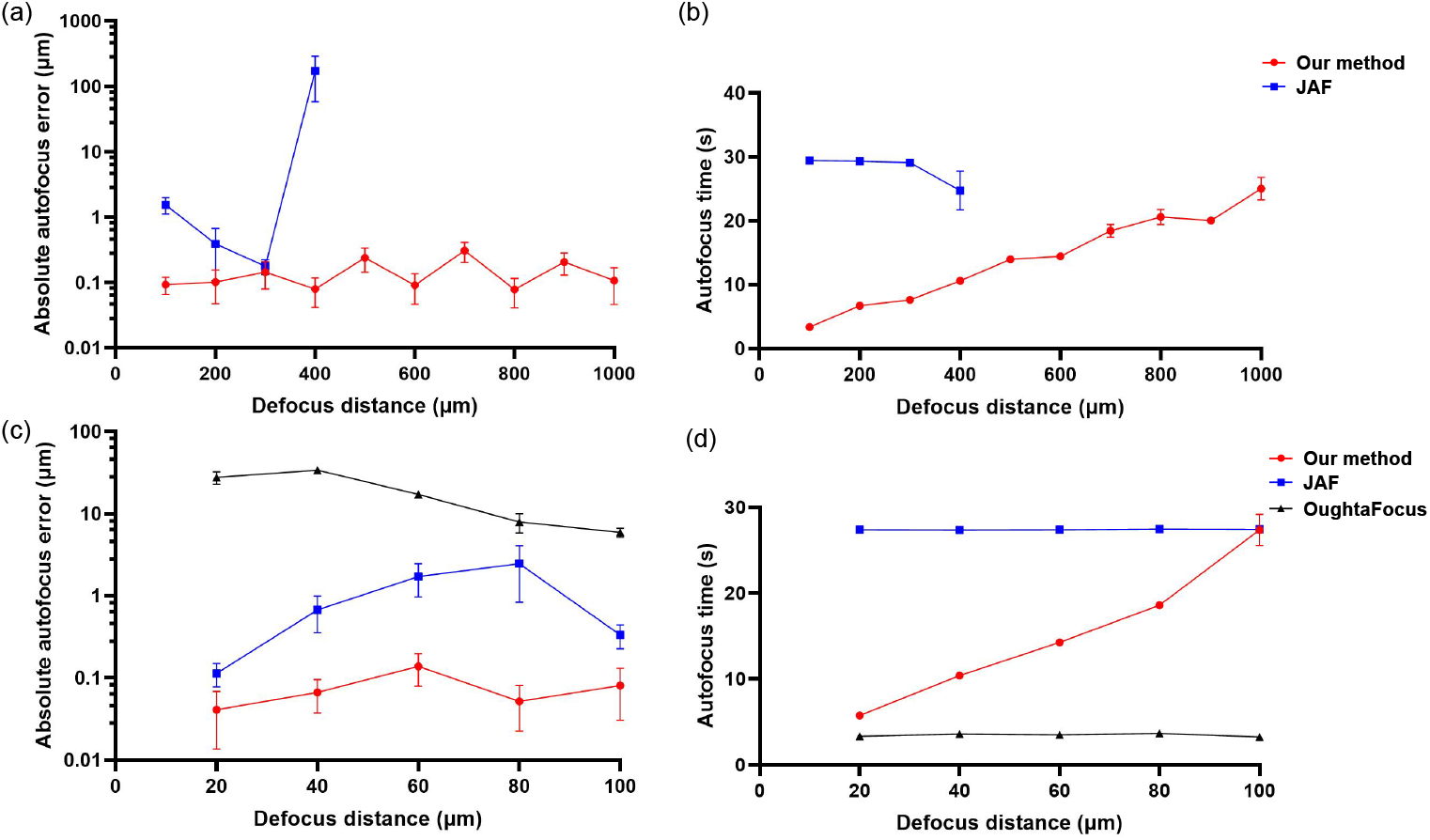
Our autofocus method outperforms JAF and OughtaFocus at high magnification in both accuracy and speed. a) Our method (red) works over a wider starting error range (within 1000 μm) than JAF (blue) (within 400 μm) and achieves a lower autofocus error (logarithmic y-axis). At defocus distances greater than 100 μm, OughtaFocus (not shown) failed to find the focus. b) Our method requires less total autofocus time than JAF. c) Within 100 μm range, OughtaFocus (black) successfully finds a focus, but with a higher error than JAF (blue) or our method (red). JAF and our method use a smaller initial search step size to enable more accurate focus at small starting errors. Our method reduces the autofocus errors by 2 to 200 times compared to the others (logarithmic y-axis). d) Our method is faster than JAF but slower than OughtaFocus, particularly at large defocus distances. For each position, autofocusing was repeated 10 times with different fields of view. SEM error bars are present but not visible where they are smaller than the datapoint marker size.

The JAF method is also based on a mountain climbing search and has already been successfully deployed for widefield microscopy to acquire images with a 20X objective. [38, 39] It supports large and small step sizes, but the search numbers for each step size must be predetermined by the user. To ensure fair comparison in terms of both autofocus speed and accuracy, we select the initial parameters for the JAF method to match those of our method (Table. S1, Supplementary 1). Specifically, we adjust the step sizes and the number of search frames at each step size to ensure consistency across both methods.

The OughtaFocus method, [40, 41] uses Brent’s algorithm, [42] a search method based on the golden section principle, to narrow the search range, and parabolic interpolation to approach the optimal position. It searches for the position with the maximum sharpness value corresponding to the predicted focal plane. The evaluation function uses “SharpEdges” and “NormalizedVariance”, corresponding to the use of the Laplacian edge operator and the Variance operator in our method. With multiple pre-autofocus tests, we chose the best parameters in (Table. S2, Supplementary 1). As the autofocus error using the “SharpEdges” function always exceeded 50 μm, it was considered to have “failed” in this test and was therefore not included in Fig. 5.

The results of this comparison show that our presented autofocus method extends the search range by more than a factor of two and also achieves superior accuracy and competitive speed. Although the OughtaFocus method showed the highest speed, its search range was ten times narrower, and its autofocus error was more than 100 times greater than that of our method. Additionally, the predefined search range in the OughtaFocus method and the pre-set stride search numbers in JAF method limited their use when the initial position range was unknown. Considering the uncertainty of initial out-of-focus distance and potential search numbers in practice, our method stands out, as it requires minimal input knowledge while providing better autofocus performance. It is also broadly applicable, as we demonstrate in the rest of our results section.

### 3.2. Our autofocus method generalises to both low- and high-magnification objectives

To demonstrate the applicability of our method to a range of hardware configurations, we next tested the performance of our method for objectives with different magnifications. To enhance the accuracy of autofocus quantification, a monolayer of fluorescent beads was used, as it provides a more straightforward and less subjective focus determination compared to cells with multiple planes that could be considered in focus. Our method was evaluated using 20X 0.45 NA and 40X 0.95 NA air objectives to test its performance in low-magnification imaging. For high-magnification imaging, the method was tested with a 60X 0.7 NA air objective, a 60X 1.4 NA oil objective, and a 100X 1.49 NA oil objective. The results are shown in (Fig. 6, Table. S3, Supplementary 1). Although different magnifications and immersion mediums affect the DOF, all autofocus mean errors in (Fig. 6) fall within the theoretical DOF of each objective. This means that our method is compatible with most standard objectives and effective across a range of magnifications, making it suitable for visualising both cellular and subcellular structures.

**Fig. 6.**
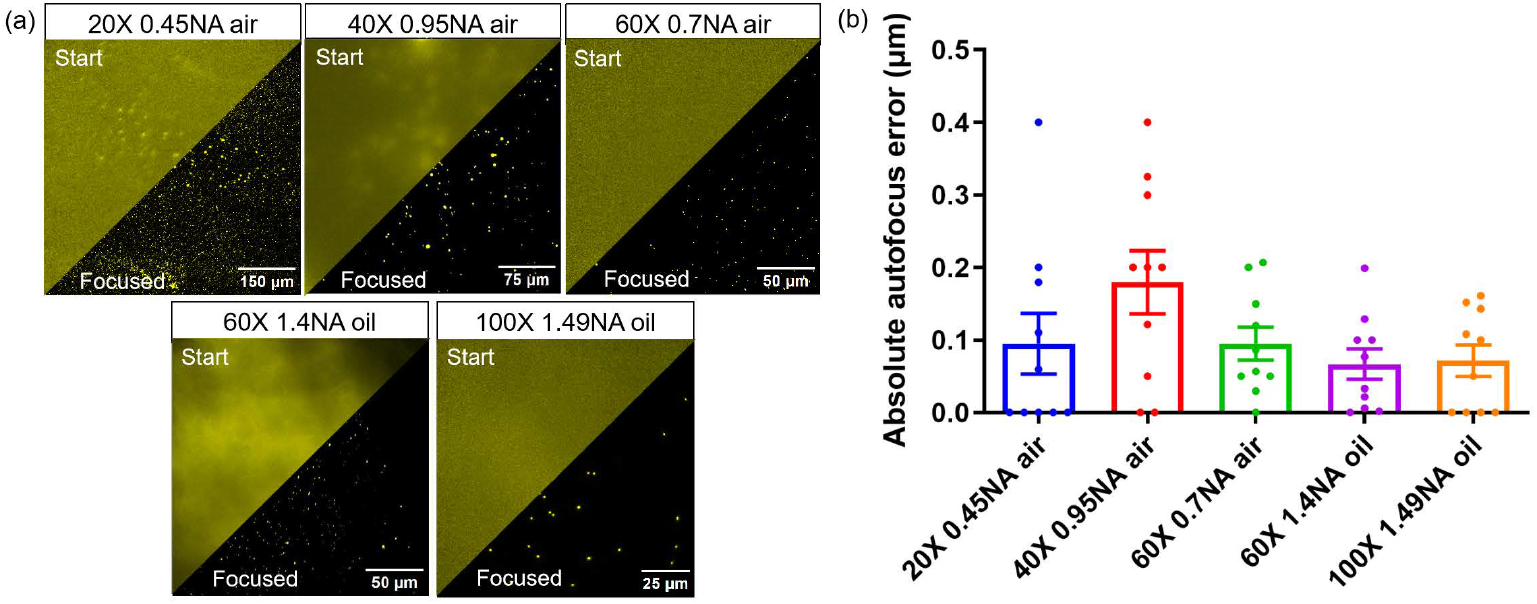
Our autofocus method performs well using objectives of different magnifications, with or without immersion oil. a) Representative fluorescence microscopy images of a sparse layer of 200 nm fluorescent microspheres before and after autofocus at magnifications from 20X to 100X. The stage’s initial start position was set to be 20 μm out of focus. b) For each objective, autofocus was repeated 10 times at different sample positions, with all the autofocus results falling within the theoretical DOF of the objectives. The results of all repeats are shown as scattered data points. The mean autofocus error with SEM is shown for each objective (see Table. S3 in Supplementary 1 for details).

### 3.3. Our autofocus method is applicable across diverse sample types and staining methods

We tested our method’s performance on a range of sample types, including fluorescent beads, and diverse subcellular structures in both live and fixed Vero and COS-7 cells. Seven subcellular structures—lysosomes, mitochondria, endoplasmic reticulum (ER), endosomes, actin, and nuclei—were chosen as they represent a range of morphologies and distributions within cells. The results are shown in (Fig. 7, Table. S4, Supplementary 1). Because bead monolayers are much thinner than most types of cells and consequently also have the highest contrast, the bead sample showed the lowest autofocus error. The autofocus error increased across different cellular structures as a result of the non-finite axial distribution of the structures (i.e. there are multiple planes that could be considered in focus). However, the mean accuracy remained within the theoretical DOF of the objective lens.

**Fig. 7.**
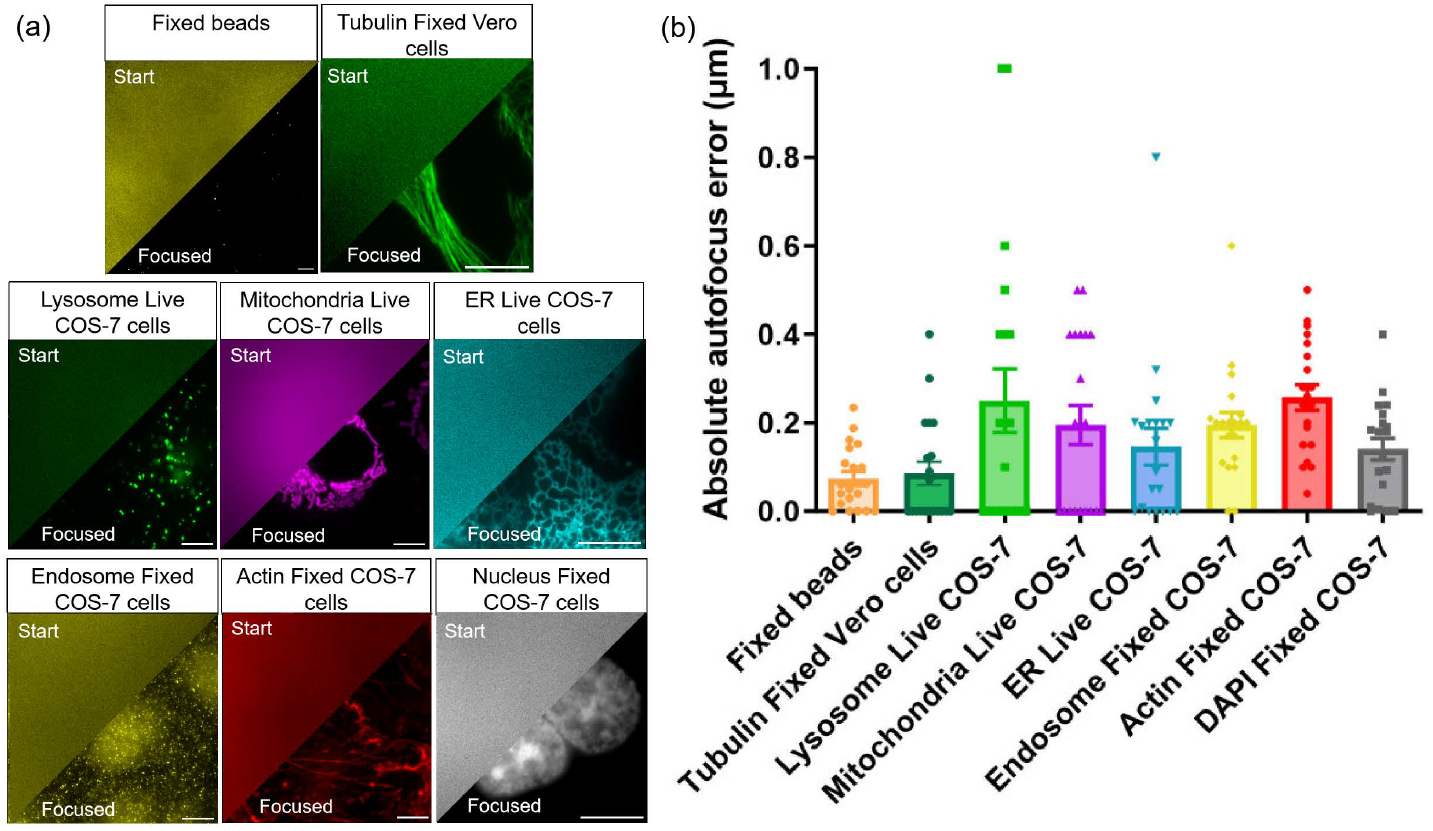
Our autofocus method performs stably well across diverse sample types and staining structures. a) Successful autofocus was achieved on various samples, including fluorescent microspheres, fixed Vero cells, and live and fixed COS-7 cells. b) Autofocus results from 20 repeats show low error rates. The mean autofocus error with SEM is plotted for each sample (see Table. S4 in Supplementary 1 for details). While live-cell imaging shows slightly less stability (larger error bars), all the results are successfully focused and reproducible. Scale bar: 10 μm. The initial stage position was set to be 20 μm out of focus, using a 100X 1.49NA objective for imaging.

### 3.4. Our autofocus method is applicable across a range of imaging modalities

In addition to epi-fluorescence widefield microscopy imaging, we further tested our method for brightfield and total internal reflection fluorescence (TIRF) microscopy (Fig. 8). In contrast to widefield fluorescence microscopy, brightfield microscopy is generally used either for imaging stained histological samples or for assessing the health of cultured cells. For some thick samples, TIRF microscopy is advantageous, as it rejects out-of-focus light, thus providing higher contrast. In widefield imaging, threshold denoising is used in our method to reduce noise in the dark background without significantly affecting the signal pattern. However, in brightfield imaging where the signal intensity is comparable to the bright background, threshold denoising will not be suitable. We therefore removed the denoising part for the brightfield microscopy imaging autofocus test. We performed a minimum of five repeats for each microscopy technique, and the method never failed, indicating its broad applicability.

**Fig. 8.**
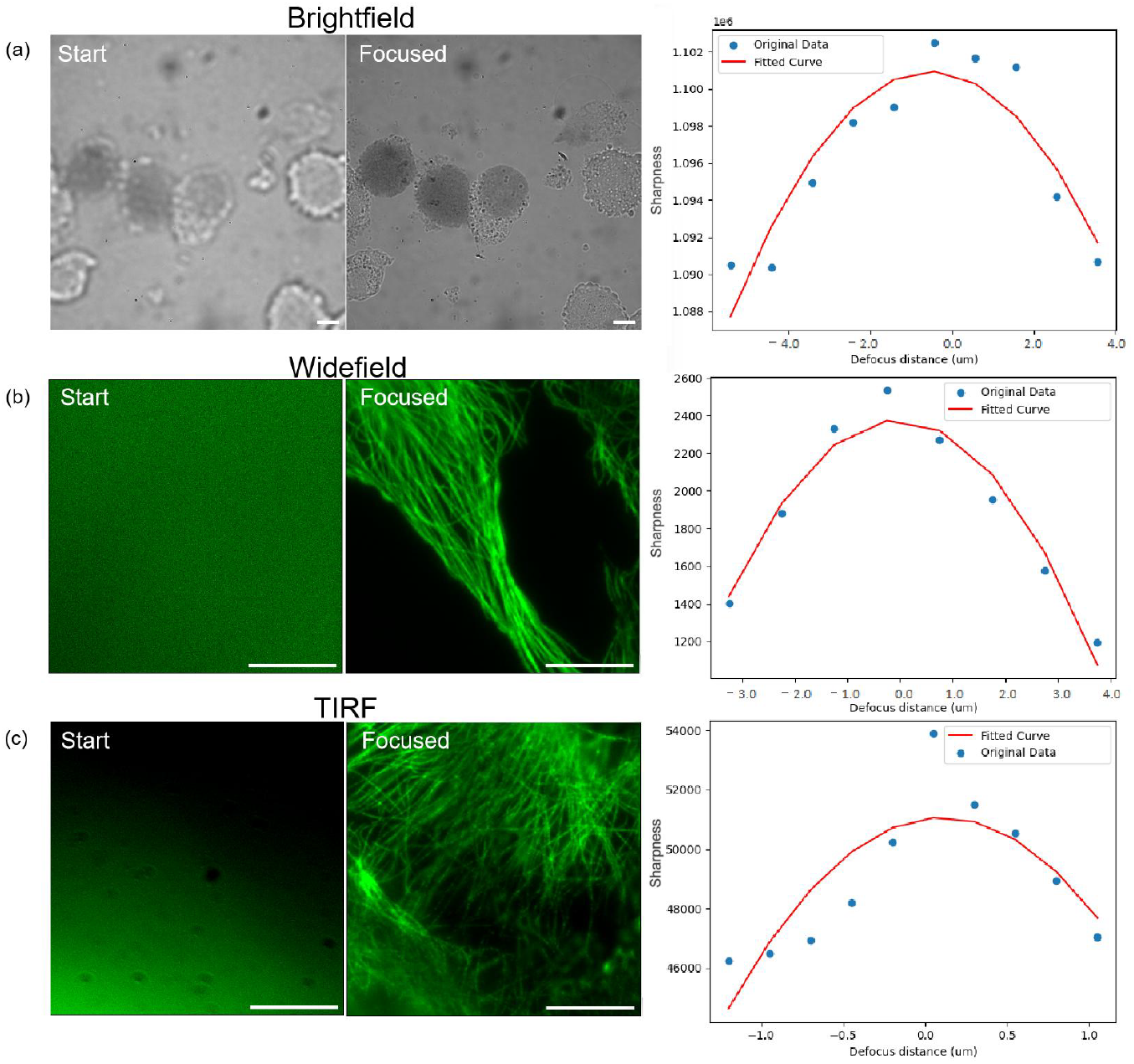
Our autofocus method is applicable to brightfield, widefield fluorescence, and TIRF microscopy techniques. The representative images show the initial start positions (left, 20 μm out of focus) and the final predicted focus positions (right), along with the fitted curves. a) The autofocus method can be applied to brightfield imaging. The sample is Trypan blue stained COS-7 cells, only dead cells appear to be with dark blue colour. b, c) Fluorescence microscopy images of immunostained *β*-Tubulin in fixed Vero cells. The autofocus method can be applied to both widefield fluorescence (b) and TIRF microscopy (c) without parameter changes.

### 3.5. Image quality improved using deconvolution

The inherent 3D nature of biological samples means that some organelles and subcellular structures are distributed across multiple planes. This not only compromises autofocus accuracy, but also causes out-of-focus light to reduce the contrast of fluorescence images. One approach to remove this out-of-focus light is to employ 3D deconvolution, where knowledge of the PSF is used to distinguish between in-focus and out-of-focus structures. With our method, this can be achieved by using the evenly spaced z-stack images acquired during autofocus. An RL-deconvolution with 20 iterations followed by maximum intensity projection of the data volume, as described above in the method section, is used to remove out-of-focus light and effectively bring structures from adjacent planes into focus simultaneously (Fig. 9).

**Fig. 9.**
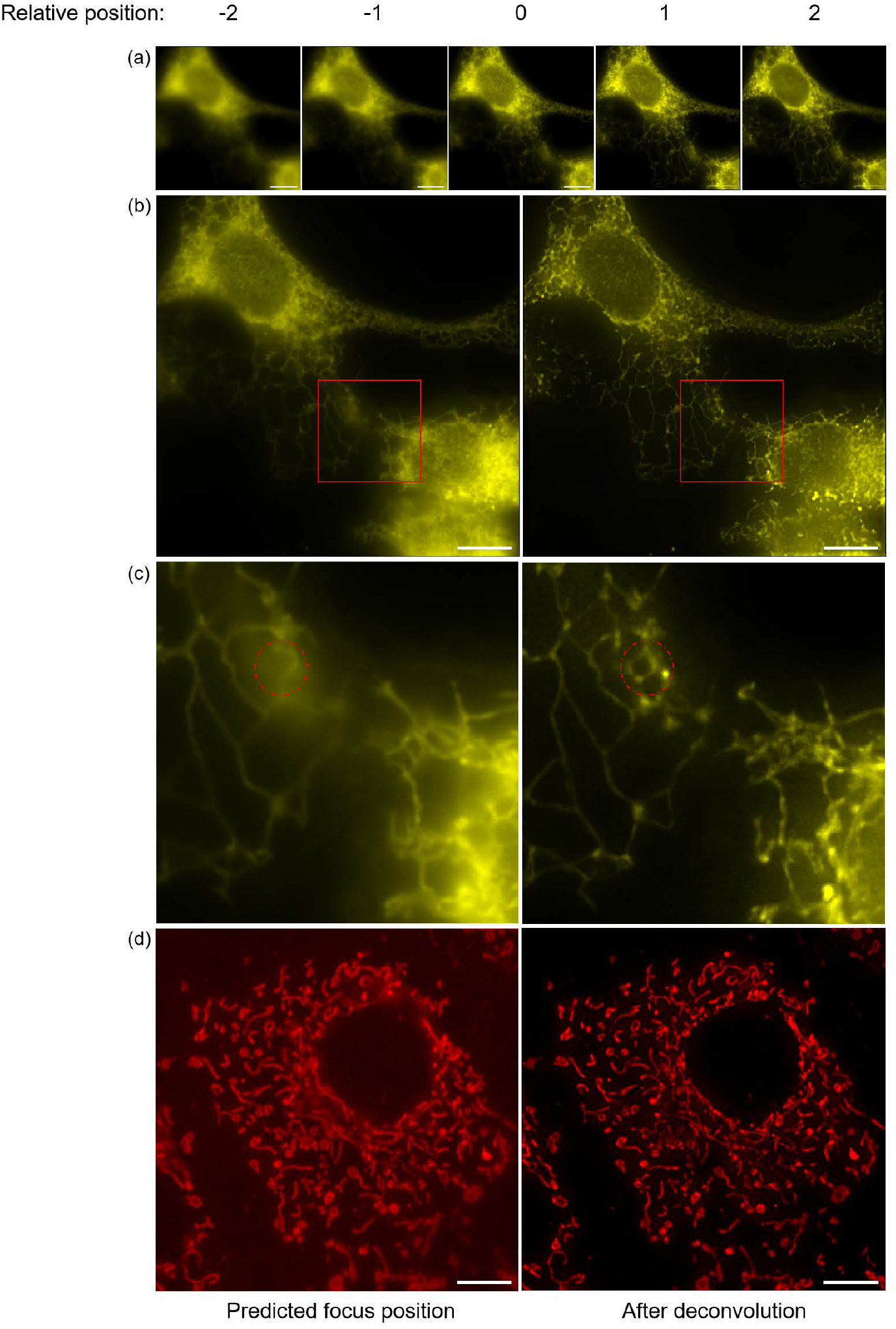
The autofocused image quality is improved by performing RL-deconvolution based on autofocus z-stacks. a) Five stack images collected from autofocus search show that different regions come into focus at different planes. The relative position indicates the negative and positive z-position of the stack (0 is the position closest to the predicted focus position), with a spacing of 500 nm between each image. b) Comparing the image at predicted focus position and after deconvolution, both the central and peripheral structures are focused simultaneously after image restoration. The sample here is live COS-7 cells with the ER labelled by VAPA-eGFP transfection. c) Magnified regions of the red rectangular areas in b). Red circles indicate structures in other planes are brought into focus after deconvolution. d) Deconvolution also improved the image quality of MitoTracker-stained mitochondria. All the images are captured with a 100X 1.49NA objective. Scale bars: 10 μm.

## 4. Discussion

In this study, we present a high performance autofocus method with broad applicability. Unlike previous methods that search with a fixed step size or rely on predefined thresholds to change the step size, our method adapts the step size using a dynamic slope threshold based on real-time analysis of the focus evaluation function. This allows automatic transition from coarse to fine search, improving the autofocus applicability, stability, and precision. It eliminates the need for repetitive searches, saving time and avoiding repeated illumination of the sample. Additionally, our search range is extended to up to 1000 μm out of focus, which means that our method is able to successfully bring the sample into focus when autonomously switching between objectives and changing FOV during biological screening.

In our method, NIQE performs well as an error-checking mechanism to ensure the precision of the final autofocus prediction, but we did not use it directly as the focus evaluation function. This is because NIQE is originally designed to evaluate the overall perceptual image quality, rather than specifically evaluating sharpness (edges and contrast), with a pretrained machine learning model using high-quality natural images. As an image goes out of focus, noise increases and gradually dominates the image. However, sometimes noise shows similar statistical features to specific natural image patterns like sand, resulting in a low NIQE score that mistakenly suggests good image quality. Therefore, using the focus evaluation function with NIQE score could be misleading because it does not have a clearly defined peak at the optimal focus. Additionally, NIQE can be more computationally intensive due to machine learning model prediction and statistical feature extraction, compared to sharpness metrics. Therefore, sharpness metrics based on Laplacian function and Variance operator are used as focus evaluation function rather than NIQE.

Although our method is primarily designed for higher-magnification imaging that requires higher autofocus accuracy, it also demonstrates competitive results at low magnification. Compared to search-based methods, such as JAF and OughtaFocus, our method reduced the autofocus errors by up to 200 times at 100X magnification. Furthermore, our method achieves significantly lower autofocus errors compared to previously reported autofocus errors in the literature at lower magnification: the autofocus error of a published mountain climbing search autofocus method [22] with adaptive search step size and curve fitting is 2.1±0.748 μm, using a 20X magnification objective on cervical cancer cells, which is more than 20 times larger than our errors (0.095±0.042 μm, Table. S3, Supplementary 1). Additionally, compared to a notable machine learning autofocus method published by Hua [14] with the autofocus error at 0.367±0.486 μm in 20X dataset, our autofocus error is still lower.

Regarding the autofocus time, which affects the throughput, our method is also competitive. It performed well on total autofocus time in direct comparison with JAF and OughtaFocus in our study. Here, we use the total autofocus time that includes the whole autofocus process from the initial position to a predicted focus position. The composition of our total autofocus time is shown in (Fig. S3, Supplementary 1). As camera capture time and autofocus calculation time are the major parts of total autofocus time, further improvement can be focused on lowering the exposure time to accelerate image acquisition and optimising the initial search step size according to specific applications. This would help to reduce the number of redundant searches and shorten the autofocus calculation time, allowing higher throughput.

The applicability of our method across a wide range of magnification objectives, diverse sample types, and three types of microscopy techniques highlights its potential for high-throughput biological screening. The screening can be combined with region of interest identification techniques at low magnification, followed by high-magnification imaging of only these key regions, further increasing the throughput and streamlining data storage. Future studies can further extend our method’s applicability to super-resolution microscopy. As super-resolution microscopy overcomes the diffraction limits, it therefore allows subtle subcellular or molecular detail visualisation, [43–45] which is especially promising in biological screening. Adapting our method for super-resolution imaging could combine the convenience and precision of autofocus techniques with the outstanding resolving capability of super-resolution microscopy techniques.

## Acknowledgment

We would like to thank Luca Mascheroni for fixed Vero cell preparation.

## Code Availability

The autofocus code and image restoration code have been uploaded to Github: https://github.com/jyt368/Autofocus

## Supplemental document

See Supplementary 1 for supporting content.

