## Supplementary 1 for "An Enhanced Mountain Climbing Search Algorithm to Enable Fast and Accurate Autofocusing in High Resolution Fluorescence Microscopy"

### 1. SUPPLEMENTARY NOTES

#### A. Image denoising

A denoising procedure was used to filter out pixels with intensities lower than a predefined threshold (set to 0), to reduce noise whilst preserving the signal within the image. In this procedure, the mean pixel intensity of the image at its initial position is calculated, then added up with its standard deviation to obtain the denoising threshold:

$$\text{Threshold} = \mu + 2 \times \sigma \tag{S1}$$

$$\mu = \frac{1}{N}\sum_{i=1}^{N} I\left(x_i, y_i\right) \tag{S2}$$

$$\sigma = \sqrt{\frac{1}{N}\sum_{i=1}^{N}\left(I\left(x_i, y_i\right) - \mu\right)^2} \tag{S3}$$

Where $\mu$ is the mean pixel value and $\sigma$ is the standard deviation of the pixels.

#### B. Quadratic sharpness distribution

The defocus induced phase difference can be calculated by [1, 2]:

$$\phi_{(x,y)} = -\frac{\Delta z \cdot n\left(x^2 + y^2\right)}{2R^2} = B\Delta z \cdot n\left(x^2 + y^2\right) \tag{S4}$$

Where $\Delta z$ indicates the defocus distance, $n$ is the refractive index of the immersion medium between the objective and the sample, and $\left(x^2 + y^2\right)$ represents the radial distance of the exit pupil from the optical axis. $R$ is the distance between sample plane and the exit pupil, and $B$ contains the constant part of the defocus phase shift.

To analyse the point spread function (PSF) with defocus aberration, we first modelled the pupil function and then calculated the PSF by computing the squared modulus of its Fourier Transform. The pupil function $P\left(x, y\right)$ is given by:

$$P\left(x, y\right) = A \cdot e^{i\phi_{(x,y)}} \tag{S5}$$

$$A = \begin{cases} 0, & \text{(x, y) lies outside the pupil} \\ \frac{1}{\sqrt{\cos\theta}}, & \text{(x, y) lies inside the pupil} \end{cases} \tag{S6}$$

Here, $A$ represent the amplitude of the pupil function, while $\phi_{(x,y)}$ is the phase shift. The intensity is 0 outside the pupil as the pupil size blocks the light from coming in, while $\frac{1}{\sqrt{cos\theta}}$ inside the pupil, where $\theta$ is the angle between incident light and the optical axis. The amplitude $\frac{1}{\sqrt{cos\theta}}$ is due to the light intensity projection at the exit pupil plane. Then, PSF can be calculated as:

$$
\begin{aligned}
\mathrm{PSF}(x', y') &= \left| \int \int P(x, y) e^{-i2\pi(f_x x + f_y y)} \, dx \, dy \right|^2 \\
&= \left| A \int_0^R \int_0^{2\pi} e^{iB\Delta z r^2} e^{-i2\pi f r \cos(\theta - \theta_f)} r \, dr \, d\theta \right|^2 \\
&= \left| A \int_0^R e^{iB\Delta z r^2} e^{2\pi J_0(2\pi f r)} \, dr \right|^2
\end{aligned} \tag{S7}
$$

Using radical coordinates, $\theta_f$ represents the angle of frequency component $(f_x, f_y)$ in the frequency domain. $J_0(2\pi f r)$ is the zero-order Bessel function deduced from Bessel integrals. [3, 4] Then, because $\Delta z \ll R$, $e^{iB\Delta z r^2}$ can be approximate to:

$$
e^{iB\Delta z r^2} \approx 1 + iB\Delta z r^2 \tag{S8}
$$

Therefore, PSF $(x', y')$ consists of a constant part and a part that proportional to $\Delta z^2$. Because Optical Transfer Function (OTF), which describes the contrast and phase changes in an image, is the Fourier Transform of PSF, OTF $(x', y')$ is also proportional to $\Delta z^2$.

#### C. Microscopy system for autofocus

We validated the performance of our autofocus method in fluorescence microscopy imaging. We used a custom-built widefield microscope based on an Olympus IX83 inverted microscope, equipped with a motorised stage and an LED light source (LED4D067, Thorlabs, Newton, New Jersey, USA) with excitation wavelengths of 405 nm, 470 nm, 590 nm, and 625 nm. The system allows both brightfield and widefield fluorescence microscopy modes. The filter cubes we used here are 39000 AT - DAPI, 39002 AT - EGFP, 39007 AT - Cy5, and 39010 AT - mCherry, all from Chroma, Bellows Falls, Vermont, USA.

We also used a custom-built Total Internal Reflection Fluorescence (TIRF) microscope with HiLo/TIRF mode to further validate our autofocus performance with better adaptability. The laser we used in this TIRF microscope is 561 nm (Omicron, LightHUB), and the laser intensity was set to 130 mW cm$^{-2}$. The filter cube we used here is FF01-600/52-25 (Semrock, Rochester, New York, USA).

### 2. SAMPLE PREPARATION

#### A. Fluorescent beads

Bead monolayer was prepared by air drying 4 $\mu$L of 0.2 $\mu$m TetraSpeck microspheres (T7280, Thermo Fisher Scientific, Inc., Waltham, MA, USA) at $10^{-5}$ dilution into an 8-well $\mu$-slide (80821, Ibidi, Gräfelfing, Germany). Wells were then filled with 500 $\mu$L of 1X phosphate buffered saline (pH 7.4).

#### B. Vero cells

Vero cells (derived from African green monkey kidney epithelial cells) were plated into 8-well slide dishes (Ibidi, Gräfelfing, Germany), twenty thousand cells per well, and cultured under standard conditions (37°C, 5% $CO_2$) in minimum essential medium (Sigma Aldrich h, Inc., St. Louis, MO, USA) supplemented with 10% foetal bovine serum (FBS, Gibco, CA, USA) and 2 mM L-glutamine (GlutaMAX, Gibco, CA, USA). After 24 h, cells were fixed by incubation with 4% methanol-free formaldehyde and 0.1% glutaraldehyde in cacodylate buffer (pH 7.4) for 15 min at room temperature, washed three times with PBS and then permeabilised by incubation with a 0.2% solution of saponin in PBS for 15 min. Unspecific binding was blocked by incubating with 10% goat serum and 100 mM glycine in PBS and 0.2% saponin for 30 min at room temperature. Without washing, the samples were incubated with the primary mouse anti-beta-tubulin antibody (Ab131205, Abcam, Cambridge, UK) diluted 1:200 in PBS containing 2% bovine serum albumin (BSA) and 0.2% saponin overnight at 4°C. After three washes in PBS, the samples were incubated with the secondary goat anti-mouse antibody conjugated to AlexaFluor568 (A12380, Thermo Fisher Scientific, Inc., Waltham, MA, USA) diluted 1:400 in PBS containing 2% BSA and 0.2% saponin for 1 h at room temperature in the dark. Samples were then washed 3 times with PBS and imaged.

### C. COS-7 cells

#### *C.1. Cell culture*

COS-7 cells (African green monkey kidney fibroblast-like cells) from the American Type Cell Collection (ATCC, Manassas, VA, USA) were grown in T25 flasks (Greiner Bio-One, Austria) by incubation at 37°C in a 5% $CO_2$ atmosphere. Complete medium for normal cell growth consisted of 90% Dulbecco's modified Eagle's medium (DMEM), 10% FBS, and 1% streptomycin. Splitting was performed at >80% confluency, and medium was refreshed every 3–4 days. For imaging, cells were cultured in 8-well slide dishes (Ibidi, Gräfelfing, Germany).

#### *C.2. Preparation of fixed samples*

Cells were fixed for 15 min using 4% PFA in PBS at room temperature. Cells were then washed 3 times for 5 min using PBS. Permeabilisation was carried out for 10 min using PBS-T (0.1% Triton in PBS). Cells were again washed 3 times for 5 min using PBS. Cells were then blocked for 1h at room temperature using 5% donkey serum. Primary staining was then performed for 2 h at room temperature. A rabbit anti-Rab5 antibody (ab13253, Abcam, Cambridge, UK) was used. Following primary staining, two quick washes using PBS were performed, followed by five 3-minute washing steps. Secondary staining was then performed for 1h using a donkey anti-rabbit antibody conjugated to Alexa Fluor 488 (A-21206, Thermo Fisher Scientific, Inc., Waltham, MA, USA) at a dilution of 1:2000 in PBS. During secondary staining, direct staining of filamentous actin was performed using Phalloidin-Atto7647 (65906, Merck, Darmstadt, Germany) at a dilution of 1:40 in PBS. Two quick washes using PBS were then performed. Cells were subsequently incubated with 1:1000 DAPI for 15 min at room temperature. Finally, cells were washed 5 times for 3 min using PBS.

#### *C.3. Live imaging*

For ER labelling, cells were transfected with plasmid pEGFPC1-hVAP-A (encoding human VAP-A fused to GFP) with Lipofectamine 2000 (Invitrogen, Inc., Paisley, UK) according to the manufacturer's protocol 24 hours before imaging. 200 ng of plasmid was used per well. pEGFPC1-hVAP-A was a gift from Catherine Tomasetto (104447, Addgene, Watertown, USA). Cells were imaged in a microscope stage top micro-incubator (OKO Lab, Pozzuoli, NA, Italy) with continuous air supply (37°C, 5% $CO_2$).

For lysosome staining, cells were incubated with 0.3 mg/mL Dextran, Alexa Fluor 594 (D22913, Invitrogen, Thermo Fisher Scientific, Inc., Waltham, MA, USA) for 4 hours, followed by washing once with culture medium, and incubated with fresh culture media for 20 hours.

For mitochondrion staining, cells were stained with 50 nM MitoTracker Deep Red (M22426, Invitrogen, Thermo Fisher Scientific, Inc., Waltham, MA, USA) for 30 minutes. They were then washed once with culture medium and subsequently imaged at once.

#### *C.4. Preparation of dead cells for brightfield imaging*

For trypan blue stained dead cells, a trypsinised cell suspension was transferred to a 15 mL centrifuge tube and kept at room temperature for two hours to accelerate the cell death process. Then, 10 $\mu$L of cell suspension was diluted with 0.4% Trypan Blue Stain (Cat.15250061, Gibco, CA, USA) at a ratio of 1:1. This mixture was kept at room temperature for 5 minutes and transferred to cover glass slide for brightfield imaging.

### 3. SUPPLEMENTARY TABLES

**Table S1. Parameters for JAF autofocus tests. Large range means the initial out-of-focus position is between 100-1000 $\mu$m (where 0 is the optimal focus plane). Small range means the initial out-of-focus position is within 100 $\mu$m.**

| JAF | Small range | Large range |
|---|---|---|
| Large step size ($\mu$m) | 1 | 10 |
| Large step number | 50 | 50 |
| Small step size ($\mu$m) | 0.5 | 5 |
| Small step number | 10 | 10 |
| Threshold | 0.02 | 0.02 |
| Crop ratio | 1 | 1 |

**Table S2. Parameters for OughtaFocus autofocus tests. Small range means the initial out-of-focus position is within 100 $\mu$m.**

| OughtaFocus | Small range |
|---|---|
| Search range ($\mu$m) | 250 |
| Tolerance ($\mu$m) | 0.1 |
| CropFactor | 1 |
| Exposure (ms) | 100 |
| FFT Lower Cutoff (%) | 2.5 |
| FFT Upper Cutoff (%) | 14 |
| ShowImages | Yes |
| Maximize | NormalizedVariance |
| Channel | DIC |

**Table S3.** **Mean autofocus errors and SEM values for different magnification objectives. Errors were quantified by comparing the algorithm-predicted focus position with the optimal focus position (Method 2.7). All the predicted focus positions fall within the theoretical depth of field.**

| | **20X 0.45NA air** | **40X 0.95NA air** | **60X 0.7NA air** | **60X 1.4NA oil** | **100X 1.49NA oil** |
|---|---|---|---|---|---|
| Repeat times | 10 | 10 | 10 | 10 | 10 |
| Mean autofocus error ($\mu$m) | 0.095 | 0.180 | 0.095 | 0.067 | 0.071 |
| Standard error of mean ($\mu$m) | 0.042 | 0.044 | 0.023 | 0.021 | 0.022 |

**Table S4.** **Mean autofocus errors and SEM values for different samples, including live and fixed samples. Live samples showed greater variations compared to the fixed samples.**

| | **Fixed beads** | **Tubulin Fixed Vero cells** | **Lysosome Live COS-7** | **Mitochondria Live COS-7** | **ER Live COS-7** | **Endosome Fixed COS-7** | **Actin Fixed COS-7** | **Nucleus Fixed COS-7** |
|---|---|---|---|---|---|---|---|---|
| **Repeat times** | 20 | 20 | 20 | 20 | 20 | 20 | 20 | 20 |
| **Mean autofocus error** ($\mu m$) | 0.074 | 0.085 | 0.250 | 0.195 | 0.146 | 0.195 | 0.257 | 0.141 |
| **Standard error of mean** ($\mu m$) | 0.016 | 0.027 | 0.072 | 0.044 | 0.042 | 0.029 | 0.029 | 0.025 |

### 4. SUPPLEMENTARY FIGURES

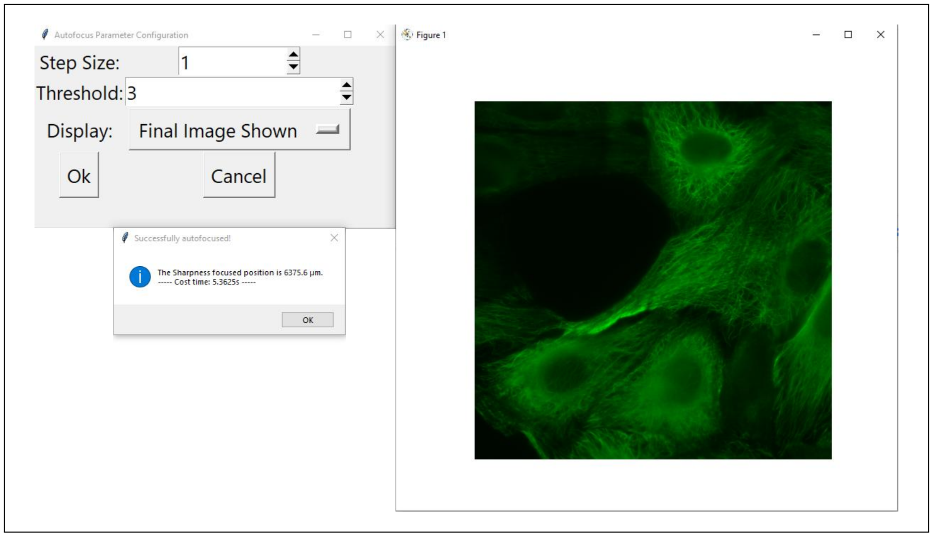


**Fig. S1.** The Micromanager-compatible autofocus interface and its output, including the final focused image, focus time, and predicted focus position. Left: The panel in which the user sets relevant parameters: initial search step size and the slope threshold. The predicted focus position and the total autofocus time are printed out once the autofocus is finished. Right: Focused image at the predicted position.

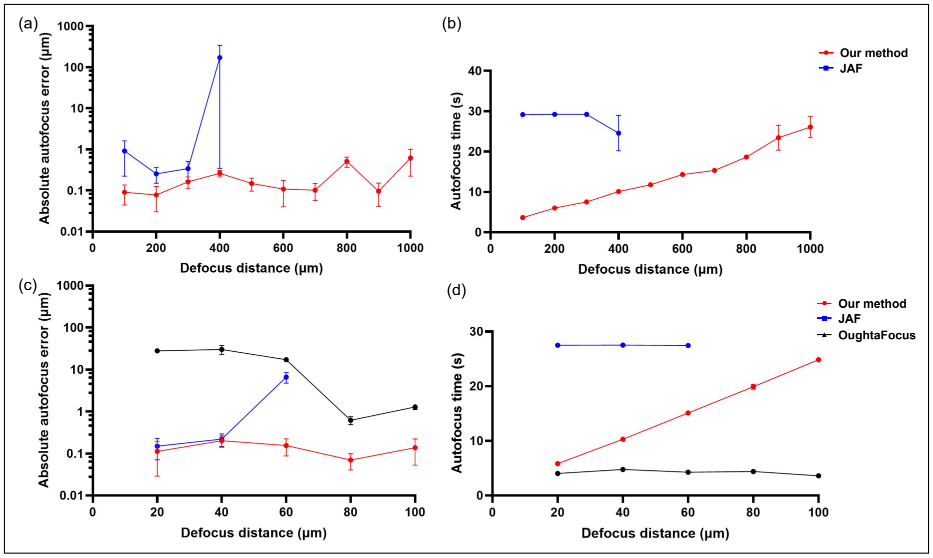


**Fig. S2.** Our autofocus method outperforms JAF and OughtaFocus in both accuracy and speed on live mitochondrion-stained COS-7 cells. a) Our method (red) works over a wider starting error range (within 1000 $\mu$m) than JAF (blue) (within 400 $\mu$m) and achieves a lower autofocus error (logarithmic y-axis). At defocus distances greater than 100 $\mu$m, OughtaFocus (not shown) failed to find the focus. b) Our method requires less total autofocus time than JAF. c) For a smaller search range within 100 $\mu$m, OughtaFocus (black) successfully finds a focus, but with a higher error than JAF (blue) or our method (red). JAF and our method use a smaller initial search stride to enable more accurate focus at small starting errors. Our method reduced the autofocus errors up to 200 times compared to the others (logarithmic y-axis). d) Our method is faster than JAF but slower than OughtaFocus. For each position, autofocusing was repeated 10 times. SEM error bars are present but not visible where they are smaller than the datapoint marker size.

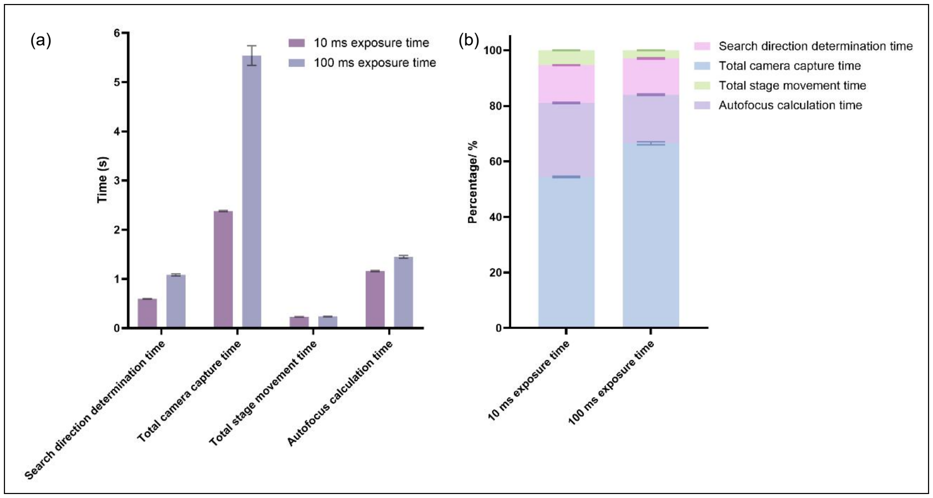


**Fig. S3.** Camera capture time and autofocus calculation time are the main contributions to the total autofocus time per field of view. a) Reducing camera capture time shortens the total autofocus time. When the exposure time was reduced from 100 ms to 10 ms, the total autofocus time was halved, with a slight reduction in autofocus calculation time. b) Camera capture time accounts for the highest percentage (approximately 50% ) of the total time, while the autofocus calculation time is the second major contributor, accounting for about 20% of the total autofocus time per field of view.
